# The roles of history, chance, and natural selection in the evolution of antibiotic resistance

**DOI:** 10.1101/2020.07.22.216465

**Authors:** Alfonso Santos-Lopez, Christopher W. Marshall, Allison L. Welp, Caroline Turner, Javier Rasero, Vaughn S. Cooper

## Abstract

History, chance, and selection are the fundamental factors that drive and constrain evolution. We designed evolution experiments to disentangle and quantify effects of these forces on the evolution of antibiotic resistance. History was established by prior antibiotic selection of the pathogen *Acinetobacter baumannii* in both structured and unstructured environments, selection occurred in increasing concentrations of new antibiotics, and chance differences arose as random mutations among replicate populations. The effects of history were reduced by increasingly strong selection in new drugs, but not erased, at times producing important contingencies. Selection in structured environments constrained resistance to new drugs and led to frequent loss of resistance to the initial drug. This research demonstrates that despite strong selective pressures of antibiotics leading to genetic parallelism, history can etch potential vulnerabilities to orthogonal drugs.

## Introduction

Evolution can be propelled by natural selection, it can wander with the chance effects of mutation and genetic drift, and it can be constrained by history, whereby past events limit or even potentiate the future (*1–5*). The relative roles of these forces has been debated, with the constraints of history the most contentious (*6*). A wealth of recent research has shown that evolution can be surprisingly repeatable when selection is strong even among distantly related lineages or in different environments (*7, 8*), but disparate outcomes become more likely as the footprint of history (*i.e*. differences in genetic background caused by chance and selection in different environments) increases (*6*) (For a detailed definition of the forces, see **Box 1**). In the absence of chance and history, selection will cause the most fit genotype to fix in the particular environment, and provided this variant is available, evolution will be perfectly predictable (*7, 9*). However historical and stochastic processes inevitably produce some degree of contingency, making evolution less predictable, reflecting the importance of evolutionary history (*3, 6, 10, 11*). The evolution of a new trait, whether by horizontally acquired genes or *de novo* mutation, is a stochastic process that depends on available genetic variation capable of producing a new trait (*12, 13*).

As any other evolved trait, antimicrobial resistance (AMR) is subject to these three evolutionary forces (Box 1). Antibiotics can impose strong selection pressure on microbial populations, leading to highly repeatable evolutionary outcomes (*14, 15*), with the level of parallelism predicted to depend on the strength of antibiotic pressure (*16*). However, evolutionary history can also alter the distribution of fitness effects of AMR mutations, their mechanisms of action, or their degree of conferred resistance (*17*). For example, the effects of a given mutation can vary in different genetic backgrounds (epistasis) or in different environments (pleiotropy) (*17–21*). Additionally, chance differences in the mutations acquired, their order of occurrence, or compensatory mutations that decrease resistance costs can affect the eventual level of resistance and its evolutionary success in the population (*12, 16*).

### Box 1. Definitions of selection, chance and history in the evolution of AMR.

Antibiotics impose strong selective pressures on microbial populations, which can produce highly repeatable outcomes when bacterial population sizes are large and mutations are not limiting. In the absence of chance and history, **selection**, the process by which heritable traits that increase survival and reproduction rise in population frequency, will cause the fixation of the resistant allele associated with the highest fitness in the population, making evolution highly predictable. However, the origin of genetic variation is a stochastic process. **Chance** effects of acquiring a mutation, gene, or mobile element, or changes in the frequencies of these alleles by genetic drift determine whether, by what mechanism, and to what degree, resistance evolves in a given population. Further, the evolutionary **history** of a population can produces contingencies that can make evolution unpredictable. For instance, different genetic backgrounds shaped in different environments can alter the phenotype of a given mutation. History can therefore alter the occurrence, mechanism, degree, and success of antimicrobial resistance.

The study of mutational pathways to AMR has become accessible by applying population-wide whole genome sequencing (WGS) to experimentally evolved populations. Growth in antibiotics will select for resistant phenotypes whose genotypes can be determined by WGS, and their frequencies and trajectories indicate relative genotype fitness. When large populations, 1 × 10^7^ CFU/mL or higher, of bacteria are propagated, the probability that every base pair is mutated at least once approaches 99% after ~80 generations (*21*). Yet chance still remains important because most mutations are initially rare and subject to genetic drift until they reach a critical frequency of establishment, when selection dominates their fate (*22, 23*). Further, many mutations arise concurrently and those with higher fitness tend to exclude contending alleles, known as clonal interference. Thus, the success of new mutations will be determined by their survival of drift, the chance that they co-occur with other fit mutants, and by their relative fitness, which is shaped by selection and history (*24*).

The contributions of history, chance, and selection to evolution can be measured using an elegant experimental design (depicted in Figure 1A and described in detail in the Supplemental text) introduced by Travisano and coworkers (*1*), in which replicate populations are propagated from multiple ancestral strains with different evolutionary histories. This experimental design has been used to quantify effects of these forces and to predict evolution in prokaryotes, eukaryotes and even digital organisms (*1–5*), but has not been applied to study the evolution of AMR, one of the most critical threats in modern medicine. Here we use this framework to measure the relative roles of history, chance and selection in the evolution of AMR phenotypes and genotypes in the ESKAPE pathogen *Acinetobacter baumannii*, a leading agent of multidrug resistant infections worldwide and named as an urgent threat by the CDC (*25*). Quantifying contributions of these evolutionary forces is essential if we are ever to predict the evolution of drug resistance of pathogens, including microbes, HIV and malaria, and of various cancers (*26–29*).

**Figure 1.**
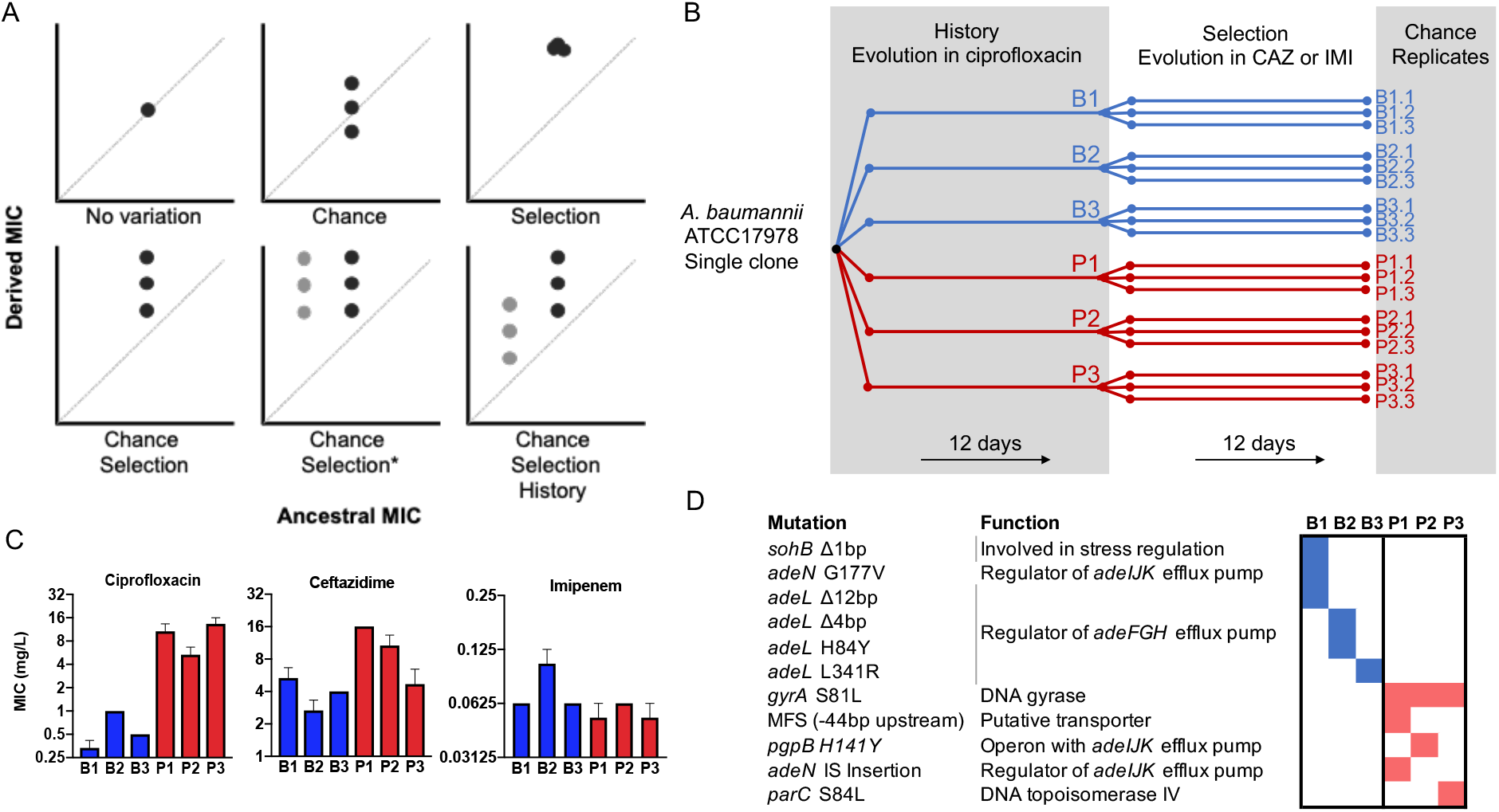
Experimental design to differentiate history, chance and selection including starting genotypes and AMR phenotypes. A) Potential outcomes of replicate evolved populations estimated by the resistance level before and after the antibiotic treatment. Grey and black symbols denote starting clones with different resistance levels. A more detailed description of this design is in the supplemental material, modified from (*1*). The asterisk denotes the case in which chance and selection both erase historical effects. B) Six different clones with distinct genotypes and CIP susceptibility were used to found new replicate populations that evolved in increasing CAZ or IMI for 12 days (*21*). C) MIC of the 6 ancestors in CIP, CAZ and IMI (+/- SEM). D) Ancestral genotypes prior to the selection phase.

## Results

Previously (*21*), we propagated a single clone of *A. baumannii* (strain 17978-mff) for 12 days in increasing concentrations of the fluoroquinolone antibiotic ciprofloxacin (CIP). In that experiment, which established the history for the present study and is analogous to prior exposure in a clinical setting, three replicate populations each were propagated in biofilm conditions or planktonic conditions (hereafter B1-B3 and P1-P3 respectively, Figure 1B). These environments selected for different genetic pathways to CIP resistance and replicate populations also diverged by chance, which produced the genetic and phenotypic histories of the ancestral strains in the current study (Figures 1C, 1D and S1). Key historical differences include reduced ceftazidime (CAZ) resistance in B populations but increased CAZ resistance in P populations (Figure 1C) (*21*).

In the current study, the “selection” phase (Figure 1B) involved experimental evolution in increasing concentrations of the cephalosporin CAZ or the carbapenem imipenem (IMI) for 12 days via serial dilution of planktonic cultures. CAZ or IMI concentrations were doubled every three days (*ca*. 20 generations), starting with 0.5X each individual clone minimum inhibitory concentration (MIC, Table S1) and finishing with 4X MIC, exposing each population to the same selective pressure during the evolutionary rescues. In this study design (Figure 1A, Supplementary Text), the extent of increased resistance represents selection, the contributions of chance are the variation among triplicate populations propagated from the same ancestor, and differences between populations derived from different ancestors quantifies effects of history (Figure 1B). The genetic effects of chance, history and selection were also determined by sequencing whole populations to a mean site coverage of 358 ± 106 bases at the end of the experiment.

### Contributions of evolutionary forces under antibiotic treatment

Antibiotic treatments usually target advanced infections, which implies large bacterial population sizes (10^8^-10^9^ cells (*30*)). Estimates suggest that a typical antibiotic treatment above the MIC concentration will clear the infection with a probability higher than 99% (*31*). But some bacterial infections can be established from as few as 10 cells (*32*), so if a few hundred cells survived the treatment this surviving subpopulation could re-infect the host. Thus, we might expect that strong selection imposed by antibiotics acting on large populations would be powerful enough to overwhelm the constraints of history. The large population sizes also might mean that many mutations are accessible in each infection, which would diminish the effects of chance. However, the bottleneck produced by the antibiotic could increase effects of drift and amplify contributions of chance and history. By propagating large populations under sequential bottlenecks, we can reproduce some of the population dynamics of the establishment and clearance of infections, and by applying Travisano et al.’s framework (*1*) we can quantify the roles of history, chance and selection in adaptation to antibiotics.

As drug concentrations increase, the strength of selection relative to other forces is also expected to increase. We therefore analyzed resistance phenotypes after 3 days of evolution under subinhibitory drug concentration and after 12 days of evolution in increasing drug levels that concluded at four times the MIC. After three days of growth in subinhibitory concentrations of CAZ, history explained the largest variation in resistance phenotypes (61.7% of variation, p<0.05), with 30.7% for selection and only 7.6% chance (Figures 2A and 2E, Methods). As expected, CAZ resistance increased overall, but some individual populations did not differ significantly from their ancestor (populations P2, P3, Figure 2A). By day 12, following propagation in 4x MIC CAZ, the amount of variation explained by selection increased to 47.8%, while effects of history dropped to 31.4% (Figures 2B and 2E), indicating that strong selective pressures can diminish or erase the effects of history.

**Figure 2.**
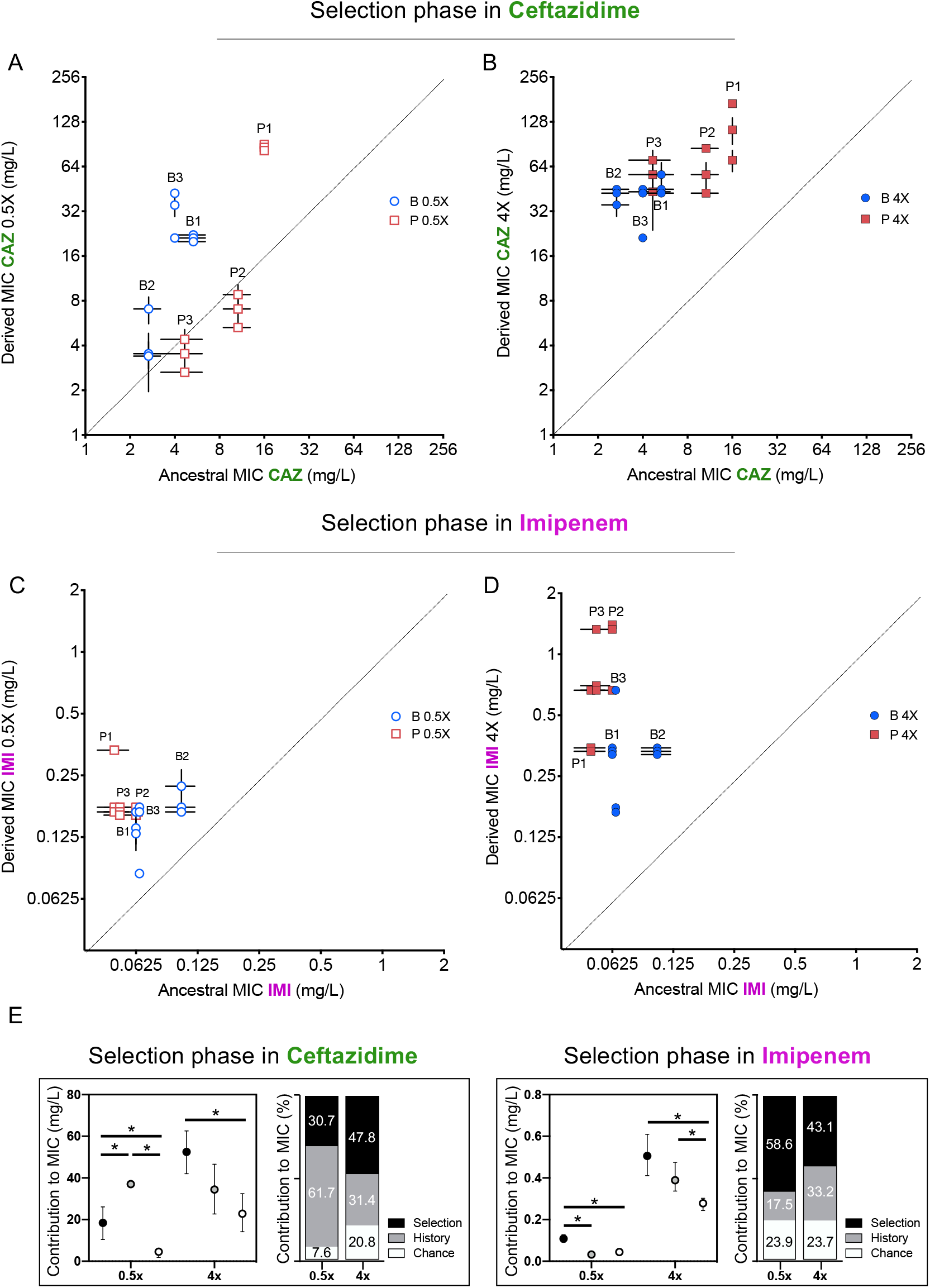
Effects of history, chance, and selection on the evolution of CAZ or IMI resistance after 3 days at 0.5x MIC (A and C) and after 12 days of increasing concentrations (B and D). Empty and filled symbols (3 days, left; and 12 days, right) represent CAZ or IMI MIC after 3 and 12 days of evolution. Blue symbols evolved from B ancestors were isolated from prior biofilm selection; red squares were evolved from P ancestors with a prior history in planktonic culture. Some symbols representing identical data points are jittered to be visible. MICs were measured in triplicate and shown +/- SEM. All populations increased CAZ resistance at day 3 (nested 1-way ANOVA, Tukey’s multiple comparison tests MIC day 0 vs. MIC day 3, p = 0.0080 q = 4.428, df = 51), and at the end of the experiment (nested 1-way ANOVA Tukey’s multiple comparison tests MIC 0 vs. MIC day 12, p = < 0.0001, q = 11.12, df = 51). All populations increased IMI resistance at day 12 but not at early timepoints (day 3) (nested 1-way ANOVA Tukey’s multiple comparison tests MIC at day 0 vs. MIC at day 12, p < 0.0001, q = 9.519, df = 51; MIC at day 0 vs. MIC at day 3, p = 0.3524, q = 1.969, df = 51). E) Absolute and relative contributions of each evolutionary force. Error bars indicate 95% confidence intervals. Asterisks denote p < 0.05.

Previous studies have shown that other evolved traits such as fitness itself show declining adaptability: less fit populations adapt faster and to a greater extent than more fit populations when propagated under the same environmental conditions (*4, 33*), which would lead to reduced variance in fitness traits among populations. This homogeneity indeed emerged as prolonged CAZ selection overcame historical variation. Populations with lower initial MICs, which by necessity were exposed to lower concentrations of CAZ, increased their resistance level more than populations with higher MICs (Figure S3), implying weak selection for further resistance in populations exceeding the MIC threshold and hence declining rates of resistance gains. This finding also suggests that the level of evolved resistance converges and may be predictable (*3, 4*), but effects of genetic background remain (Figure 2). Strong antibiotic selection has the potential to overcome but not entirely eliminate historical differences in resistance.

### Evolutionary tradeoffs arise from past antibiotic selection

Evolutionary tradeoffs occur when changes in a given gene or trait increase fitness in one environment but reduce fitness in another. For example, a history of adaptation to one antibiotic could alter the evolutionary rate and the level of resistance in the presence of a subsequent antibiotic. The phenomena of cross-resistance and collateral sensitivity are specific examples of pleiotropy, where the mechanism of resistance to the initial drug either directly increases or decreases resistance to other drugs, respectively (*34*). Additionally, the resistance mechanism could interact with other genes or alleles in the genome, a form of epistasis, and also promote or impede resistance evolution. We hypothesized that resistance mechanisms arising during selection in CAZ would alter resistance to other antibiotics both by genotype-independent (pleiotropy) and genotype-dependent (epistasis) mechanisms. Recall that during the history phase of the experiment (*21*), populations propagated in increasing concentrations of CIP became from 4 to 200-fold more resistant to CIP (Figure 1C, (*21*). Some of these strains also became more resistant to CAZ (populations P1-P3) while others became more susceptible (populations B1 and B3, for more details see reference (*21*)), and given that these populations were founded by the same ancestor this variation in collateral resistance phenotypes is best explained by pleiotropy. In the current study, after 12 days evolving in the presence of CAZ, the grand mean of CIP resistance levels did not change, so history was the dominant force shaping the MIC to CIP (Figure 3A). However, if we analyze the P and the B populations independently, B populations became significantly more sensitive to CIP but the P populations did not (Figure 3A), showing that the emergence of collateral sensitivity may depend on prior selection in different environments. These results also indicate that CAZ resistance mechanisms interact with CIP resistance in potentially useful ways.

**Figure 3.**
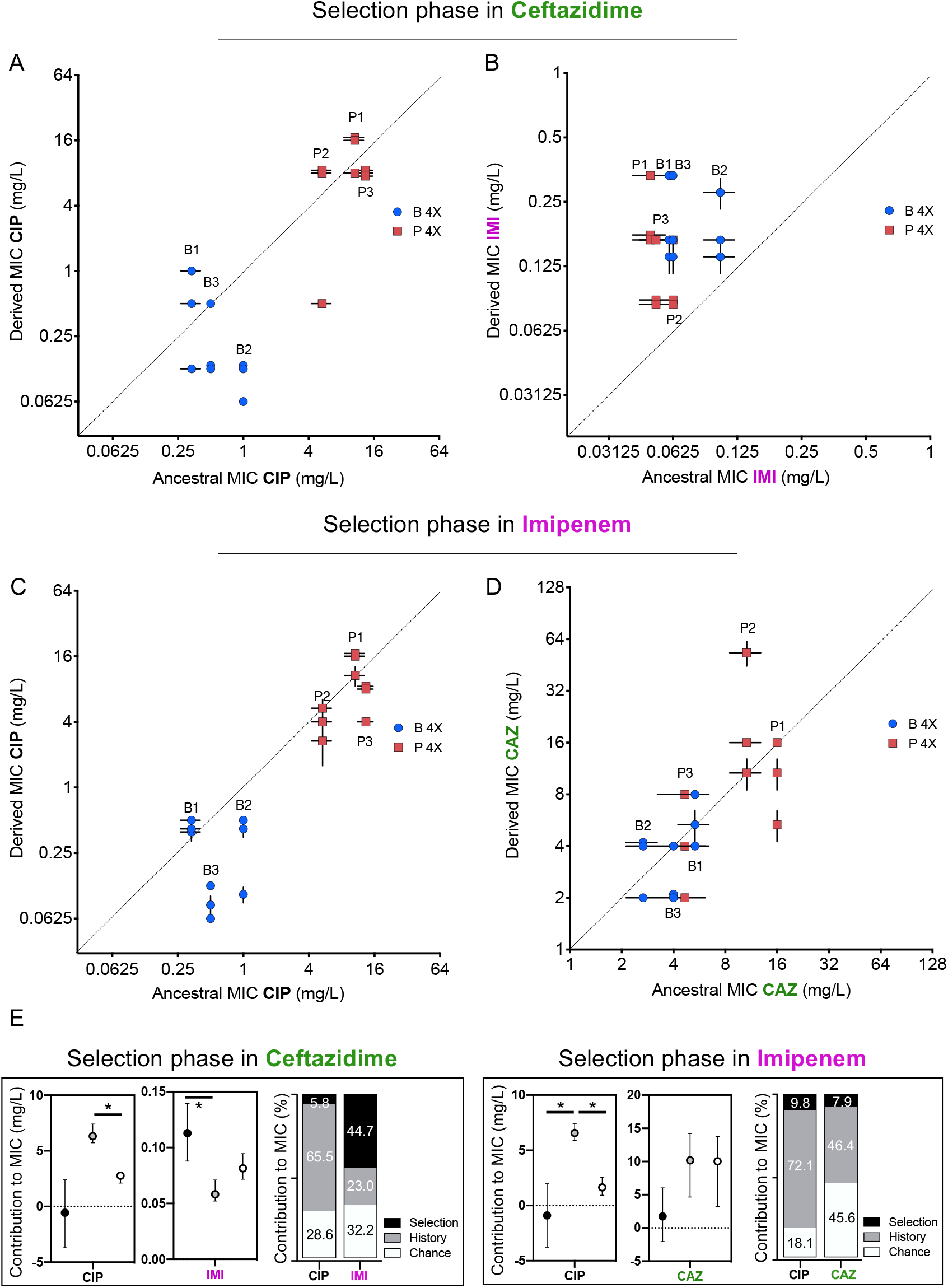
Collateral resistance caused by history, chance, and selection. Panel (A) shows CIP resistance and (B) shows IMI resistance following 12 days of CAZ treatment. Panel (C) shows CIP resistance and (D) shows CAZ resistance following 12 days of IMI treatment. Blue symbols: populations evolved from B (biofilm-evolved) ancestors; red squares: populations evolved from P ancestors (planktonic-evolved). Some symbols representing identical data points are jittered to be visible. MICs were measured in triplicate and shown +/- SEM. E) Contributions of each evolutionary force. Error bars indicate 95% confidence intervals. Asterisks denote p < 0.05.

We also tested if evolving in the presence of CAZ altered resistance to the carbapenem antibiotic IMI (Figure 3B). We hypothesized that as CAZ and IMI are both β–lactam antibiotics and mutations in efflux pumps can alter resistance to both (*35*), selection in CAZ will also increase IMI resistance and further, the contributions of each evolutionary force to IMI resistance would follow that measured for CAZ (Figure 2B). As expected, all 12 populations evolved in CAZ became more resistant to IMI (two-tailed nested t-test p < 0.0001, t = 7.507, df = 34), and selection was the most important force (p<0.05), explaining almost 44.3% of the variation, while history contributed 23.0% and chance 32.2% (Figure 3E).

### Replaying the tape of life in a different antibiotic

We learned that the evolution of resistance in *A. baumannii* to one drug, CAZ, is substantially influenced by prior history of selection in another drug, CIP, as well as the prior growth environment, planktonic (P) or biofilm (B). Namely, B-derived populations evolved CAZ resistance at the expense of their prior CIP resistance, reversing this pleiotropic tradeoff. To test if these results are repeatable and not limited to CAZ and CIP, we replayed the “selection phase” with the same genotypes using the carbapenem IMI (Figures 1, and 2 (*21*)). Here, no overall change in resistance occurred following 3 days in subinhibitory concentrations of IMI (Figure 2C) but did increase by experiment’s end at 4X MIC (Figure 2D). After the subinhibitory treatment, the more sensitive populations experienced greater gains in IMI resistance than the less sensitive populations, erasing some effects of history (Figures 2C and S3). In total, selection again predominated (p < 0.05) and explained 43.1% of the phenotypic variation in this experiment, while history explained 33.2% (Figures 2B, 2D and 2F).

As predicted by the CAZ experiment, evolution in IMI did not affect CIP resistance on average and history explained 75% of the variation in MIC (Figure 3F), but again produced collateral sensitivity in two B populations (Figure 3C). This result demonstrates that mechanisms of IMI resistance interact with historical resistance to CIP and produce tradeoffs. The biggest difference between the CAZ and IMI experiments is an asymmetry in cross-resistance between these drugs. Selection in CAZ increased IMI resistance (Figure 3B), but not *vice versa* (Figure 3D). These divergent cross-resistance networks result from the particular mutations that were selected in both experiments, which are explained below.

### Phenotypic divergence despite genetic parallelism

When multiple lineages evolve independently in the same environment, phenotypic convergence is usually observed, but the genetic causes may be more variable (*3, 4, 36*). In our experiment, large populations were exposed to strong antibiotic pressure, so we predicted parallelism at the genetic level owing to few solutions that improve both fitness and resistance (*22*). We conducted whole-population genomic sequencing of all populations at the end of the experiment to identify all contending mutations above a detection threshold of 5% and analyzed the genetic contributions of history, chance, and selection using Manhattan distance estimators (Figure 4). Specifically, selection causes new mutations to rise to detectable frequencies, history can be assessed by whether previous mutations are maintained throughout the evolution experiment, and chance is revealed as genetic variation among replicates of each ancestor. Using these metrics, we infer that evolution in CAZ was shaped more by selection than history, but the opposite was seen in IMI, and effects of chance were similar in both experiments (Figure 4).

**Figure 4.**
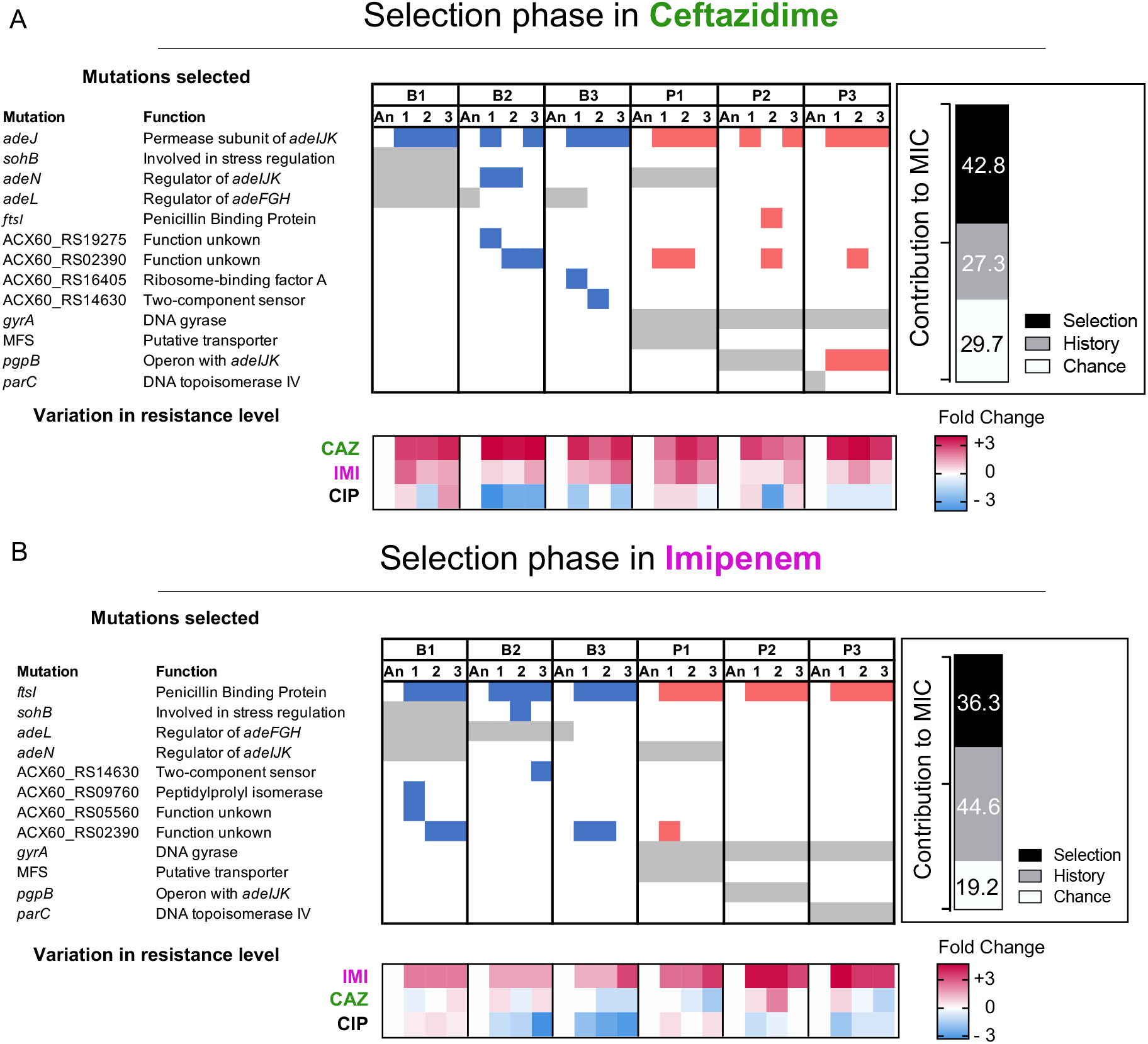
Mutated genes in the populations evolving in presence of a new antibiotic. Each column represents a population propagated in CAZ (A) or in IMI (B). Grey shading indicates the mutated genes present in the ancestral clones derived from the “history phase”. Blue and red denote mutated genes after the “selection phase” in CAZ or IMI and if those lines experienced prior planktonic selection (red) or biofilm growth (blue). Only genes in which mutations reached 75% or greater frequency or that became mutated in more than one population are shown here. A full report of all mutations is in Table S3. The relative contributions of history, chance, and selection to these genetic changes are shown in the insets. Below: log_2_ changes in evolved resistance for each population shown as a heatmap.

Clinical CAZ-resistant *A. baumannii* isolates commonly acquire mutations that increase the activity of <u>A</u>cinetobacter <u>d</u>rug <u>e</u>fflux (*ade*) pumps (*35*). In the CIP selection that established history for this study, biofilm lines (Figure 1B) selected mutations in *adeL*, the regulator of the *adeFGH* pump, which produce collateral sensitivity to CAZ and other β-lactams (Figure 1D). In contrast, P lines became cross-resistant to CAZ by *adeN* mutations that regulate the *adeIJK* complex or *pgpB* mutations that are also regulated by *adeN* (Figure 1D (*21*)). Here, evolution in increasing concentrations of CAZ selected at least one mutation in *adeJ* in 16/18 populations (Figure 4A); this gene encodes the permease subunit of AdeIJK that is a known cause of CAZ resistance (*35*). The two exception populations instead acquired mutations in *adeN*, in ACX60_RS2390, a gene of unknown function, and in *ftsI*, the target of CAZ. Evolution in IMI also selected mutations in the *ftsI* gene in all populations (Figure 4B); this gene encodes penicillin binding-protein 2, one of the most common causes of *de novo* resistance to IMI in clinical isolates (*35*). Therefore, evolution in β-lactam antibiotics generated parallel evolution regardless of the genetic background (*14, 15*).

Yet despite this genetic parallelism, replicate populations reached different resistance levels (Figures 2B and 2D). One potential reason is that different mutations in the same gene may produce different phenotypes, and another is that interactions with their genetic background – shaped by prior selection in CIP in different environments – modulate resistance levels. Evidence for the first explanation is seen when comparing replicate populations derived from ancestor P1, where different SNPs in *adeJ* (Figure 4 and Table S2) produce varied resistance (Figure 2), perhaps by altering the function of this permease. Evidence of genetic interactions can be seen when comparing the five replicate populations evolved in IMI that acquired the same mutation in *ftsI* (A579V) but differ in resistance levels by up to 4-fold owing to different historical mutations (Figure 4 and Table S2). It is currently unclear if these interactions are additive or epistatic. To summarize, both varied pleiotropy of different mutations in drug targets that balance fitness and resistance and interactions between mutations in different drug targets may therefore constrain AMR evolution.

### Collateral sensitivity resulting from genetic reversions

Antibiotic resistance mutations typically incur a fitness cost that favor sensitive strains in the absence of antibiotics. The phenotypic reversion to sensitive states is commonly caused by secondary mutations in other genes that alter resistance (*37, 38*) or it could be caused by genotypic reversions in which the ancestral allele is selected under drug-free conditions (*2, 36, 39*). In our experimental system, assuming a conservative uniform distribution of mutation rate of 10^-3^/genome/generation (*21*), each base pair experiences approximately three mutations on average during the 12 days of serial transfers (*21*). This estimate implies that reversion mutations affecting historical CIP resistance did occur, but nonetheless they are expected be much rarer than suppressor mutations in other genes. Surprisingly, we identified genetic reversion of *adeL* mutations five different times in CAZ lines and three different times in IMI lines, and these back-mutations reversed resistance tradeoffs between B-lactams and CIP (Figures 3, 4 and S2). We also observed genetic reversion of *parC* mutations in each P3 replicate propagated in CAZ (Figure 4). The topoisomerase IV *parC* is one of the canonical targets of CIP but these mutations have been shown to incur a high fitness cost in the absence of CIP (*40*). Selection therefore favored these reversions in the absence of CIP, but in this case without notable loss of CIP resistance presumably via secondary mutations in *pgpB* (Figure 4, (*21*)). The high frequency of mutational reversion observed in these experiments indicates that these resistant determinants are under enormous constraint and impose fitness costs that must be shed when the environment changes.

## Discussion

Stephen Jay Gould famously argued that replaying the tape of life is impossible because historical contingencies are ubiquitous (*41*). The evolution and spread of AMR provide a test of this hypothesis, because countless evolution experiments are initiated each day with each new prescription to combat infections caused by bacteria with different histories. Previous studies suggest that the predictability of antibiotic resistance – or the fidelity of the replay – depends on the pathogen, the antibiotic treatment, and the growth environment (*14–16, 18, 21*). Here, we have quantified contributions of history, chance and selection to AMR evolution, using six different ancestors replicated in each of two different antibiotic treatments. In the end, while selection is unsurprisingly the predominant force in the evolution of AMR, leading to parallel evolution even at the nucleotide level in some instances, history and chance play clear roles in the emergence of new resistance phenotypes (Figures 5, 3B and 3D, (*14*)), the extent of evolved resistance (Figures 2 and 3), the generation of collateral sensitivity networks (*34*)) and the predictability of the final resistance phenotype (Figures 1 and 4, *15, 18*)). Our data also suggest that, as in *Drosophila* (*39*), viruses (*36*) and yeasts (*2*), history and chance may determine the reversibility of acquired traits (Figure 5). This probability of reversion is potentially clinically important because exploitable collateral sensitivity networks can arise, such as the tradeoff between CIP resistance and beta-lactam resistance identified here (*34*). Finally, our data reveals that evolution of AMR follows a clear diminishing return pattern, where antibiotic pressure selects for mutations with progressively smaller phenotypic effects as the population is treated with higher antibiotic concentrations. This result mirrors findings in the original Travisano *et al*. paper (*1*), where populations pre-adapted to compete well in maltose did not adapt further, but populations with major deficiencies in maltose evolved to become just as fit. This result may be instructive for AMR management: on the one hand, more resistant populations at the outset did not increase this phenotype further, but on the other hand, more susceptible lines rapidly compensated for this deficit.

**Figure 5.**
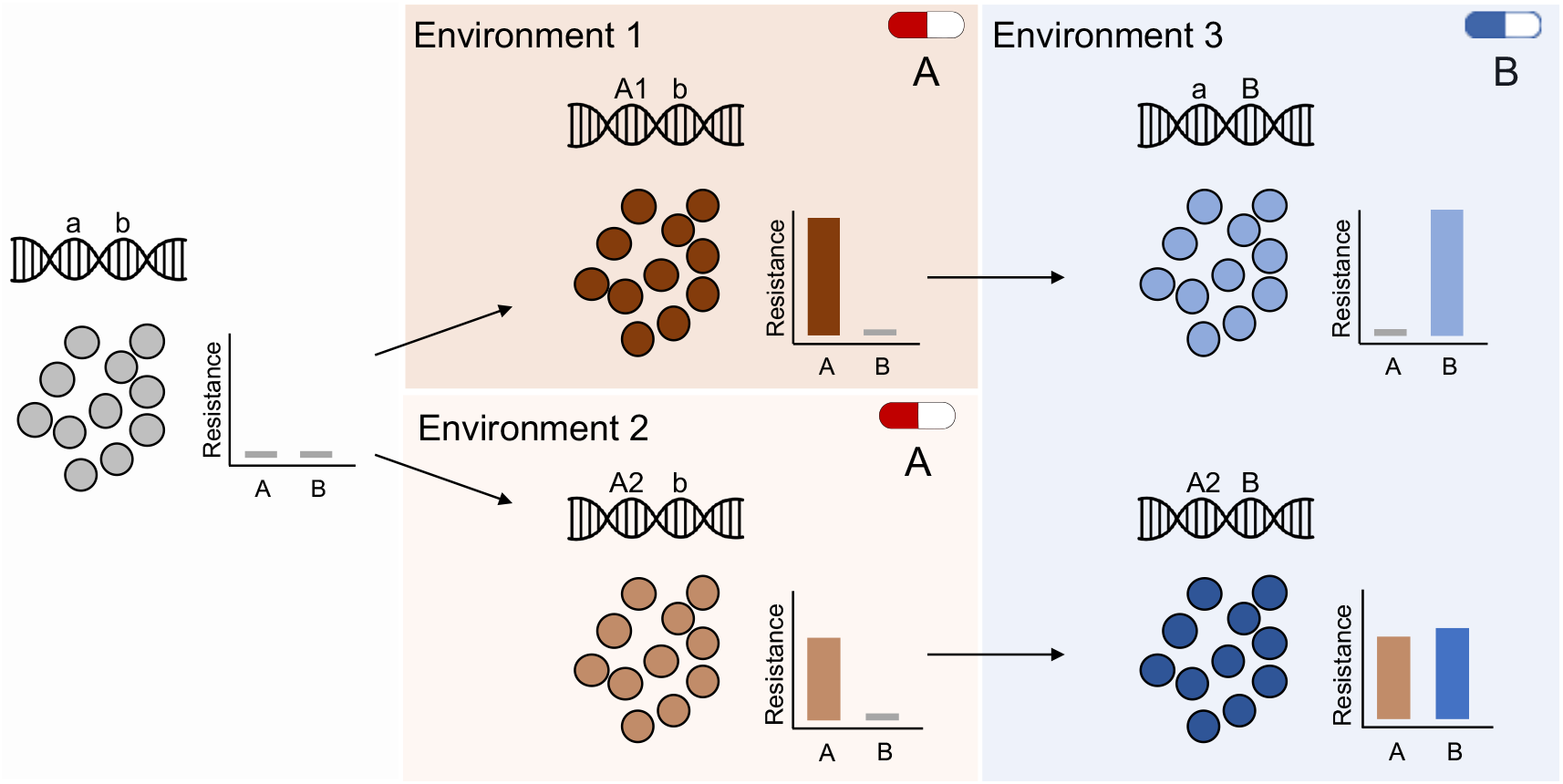
Evolutionary history and natural selection determine the evolution of antibiotic resistance. A sensitive population (left panel) is subjected to two successive treatments (antibiotic A and antibiotic B, middle and right panels respectively). First, the population was treated with antibiotic A in either of two different environments (middle panel top and bottom) that selected different genotypes (mutations A1 and A2) with distinct resistance phenotypes (middle panel insets). During subsequent exposure to a second antibiotic (B), this evolutionary history determined resistance levels (right panel) to both drugs A and B, for instance resulting in the loss of resistance to drug A (top right panel).

Our experiment focuses solely on *de novo* mutations and does not allow the opportunity for horizontal gene transfer from other species or strains, which is the principal mechanism of the emergence of antimicrobial resistances in most clinical settings (*26*). However, genetic background also affects the level of resistance conferred by transmissible elements (*42*) and epidemiological data indicate that evolutionary history constrains the persistence of resistance mediated by plasmids (*43*). The framework defined here illustrates the potential to identify genetic and environmental conditions where selection is the most dominant evolutionary force and it predictably produces antagonism between resistance traits. With ever greater knowledge of the present state, we gain hope for guiding the future to exploit the past.

## Authors contributions

VSC, AS-L and CWM conceived and designed the study; AS-L and ALW performed the experiments; CWM and CT conducted bioinformatic analysis; JR analyzed the statistical contribution of each force; AS-L and CWM, drafted the manuscript with input from all authors under VSC supervision. This work was supported by the Institute of Allergy and Infectious Diseases at the National Institutes of Health grant U01AI124302 to VSC.

## Acknowledgments

We thank Tim Cooper, Alvaro San Millan and Sergio Santos for helpful discussions and proofreading of the paper, Dan Snyder for laboratory assistance and Christopher Deitrick for depositing the sequences in the NCBI database. This research was supported by NIH U01AI124302-01.

## Methods

### Experimental evolution

#### Historical phase

Before the start of the antibiotic evolution experiment, we planktonically propagated one clone of the susceptible *A. baumannii* strain ATCC 17978-mf in a modified M9 medium (referred to as M9^+^) containing 0.37 mM CaCl_2_, 8.7 mM MgSO_4_, 42.2 mM Na_2_HPO_4_, 22 mM KH_2_PO_4_, 21.7mM NaCl, 18.7 mM NH_4_Cl and 0.2 g/L glucose and supplemented with 20 mL/L MEM essential amino acids (Gibco 11130051), 10 mL/L MEM nonessential amino acids (Gibco 11140050), and 10 mL each of trace mineral solutions A, B, and C (Corning 25021-3Cl). This preadaptation phase was conducted in the absence of antibiotics for 10 days (*ca*. 66 generations) with a dilution factor of 100 per day.

After the ten days of preadaptation to M9^+^ medium, we selected a single clone and propagated for 24 hours in M9^+^ in the absence of antibiotic. We then subcultured this population into twenty replicate populations. Ten of the populations (5 planktonic and 5 biofilm) were propagated every 24 hours in constant subinhibitory concentrations of CIP, 0.0625 mg/L, which corresponds to 0.5x the minimum inhibitory concentration (MIC). We doubled the CIP concentrations every 72 hours until 4x MIC (Figure 1B).

#### Selection phase

Upon the conclusion of the “historical phase”, we selected 1 clone from 3 populations previously adapted in biofilm and 3 populations previously adapted in planktonic conditions. We determined their resistance level to CIP, CAZ, and IMI. Then, we propagated planktonically each clone independently or in the presence of increasing concentrations of CAZ or in increasing concentrations of IMI. For each population, we used their own MIC to CAZ or IMI to determine the concentrations used in this phase (Table S1).

We froze 1mL of the control populations at days 1, 3, 4, 6, 7, 9, 10, and 12 in 9% of DMSO.

### Antimicrobial susceptibility characterization

We determined the MIC of CAZ, CIP, and IMI of the whole population by broth microdilution in M9^+^ as explained before according to the Clinical and Laboratory Standards Institute guidelines (*21*), in which each bacterial sample was tested in 2-fold-increasing concentrations of each antibiotic. The CIP, CAZ and IMI were provided by Alfa Aesar (Alfa Aesar, Wardhill, MA), Acros Organics (Across Organics, Pittsburgh, PA) and Sigma (Sigma-Aldrich Inc, St. Louis, MO) respectively.

### Genome sequencing

We sequenced the 6 ancestral clones and whole populations of the 36 evolving populations (18 evolved in the presence of CAZ and 18 evolved in the presence of IMI) at the end of the experiment. We revived each population or clone from a freezer stock in the growth conditions under which they were isolated (*i.e*. the same CAZ or IMI concentration which they were exposed to during the experiment) and grew for 24 hours. DNA was extracted using the Qiagen DNAeasy Blood and Tissue kit (Qiagen, Hiden, Germany). The sequencing library was prepared as described by Turner and colleagues (*8*) according to the protocol of Baym *et al*. (*57*), using the Illumina Nextera kit (Illumina Inc., San Diego, CA) and sequenced using an Illumina NextSeq500 at the Microbial Genome Sequencing center (https://www.migscenter.com/).

### Statistical analysis and quantification of the role each evolutionary force

We calculated the phenotypic effect of the evolutionary forces using a nested linear mixed model. By means of this nested linear mixed model including ancestors and replicates as random effects, we estimated the effect of history as the square root of the variance among all propagated populations; the effect of chance as the square root of the variance between the replicates propagated from the same ancestor; and the effect of selection was calculated as the difference in grand mean of the propagated replicates and their ancestors. (Table S4).

Percentile bootstrap was employed to compute the confidence intervals of each force at the level of significance α=0.05 by taking 1000 random samples with replacement. In addition, the statistical evidence of each force was assessed adopting a Bayesian approach, which allows to circumvent the issues associated to null hypothesis statistical testing (*45*). Specifically, a set of models excluding each force (Null hypotheses) were confronted against the full model including the three forces (Alternative Hypothesis). Thus, let BIC_1_ be the Bayesian Information Criterion associated to the alternative model and BIC_0_ the Bayesian Information Criterion for one of the null models. Then, a Bayes factor can be approximated as follows

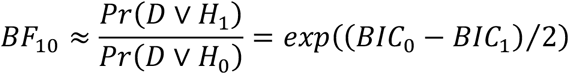

where Pr(D|H_0_) and Pr(D|H_1_) are the marginal probabilities of the data under the null and alternative models respectively. Hence, the Bayes factor allows to quantify how likely the inclusion of a force is with respect to its absence according to the observed data. All these estimations were performed using *blme* v1.0-4 R package (https://cran.r-project.org/package=blme). All values were normalized to one to calculate the influence of each evolutionary force.

The roles of the evolutionary forces at the genotypic level were calculated based on the Manhattan distance (d_M_) between populations. For a pair of populations *j* and *k* with *n* genes,

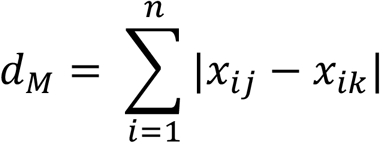

where *x_ij_* is the frequency of mutated alleles in gene i in population *j*, relative to the *A. baumannii* strain ATCC 17978-mff. The genotypic role of chance was calculated as half the mean d_M_ between all pairs of evolved populations founded from the same ancestral clone. The genotypic role of history was calculated as half the mean d_M_ between all pairs of evolved populations founded from the different ancestral clones minus the role of chance. The genotypic role of selection was calculated as the mean d_M_ between evolved populations and their founding clone, minus the roles of chance and history. In calculating selection, mutations present in the founding clone were not excluded when subtracting the effect of history.

All statistical comparisons of MIC values were performed on the log_2_ transformed values. Differences in grand means between populations were analyzed by a 1-way nested ANOVA with Tukey’s multiple comparison tests or by a nested t-test. Spearman correlation was performed using the grand means to determine the correlation between the ancestral MIC and the fold change of MIC acquired during the experiment. There are three possible outcomes by correlating the original MIC and the fold dilution change: i) a negative correlation, in which the populations with lower initial MICs increased their resistance level more than populations with higher MICs, implies that the selection erased the previous effects of history; ii) a positive correlation indicates that initial differences in MIC were magnified by selection and iii) a lack of correlation indicates that the effect of history did not change before and after selection.

### Data processing

The variants were called using the breseq software v0.31.0 (*46*) using the default parameters and the -p flag when required for identifying polymorphisms in populations after all sequences were first quality filtered and trimmed with the Trimmomatic software v0.36 (*47*) using the criteria: LEADING:20 TRAILING:20 SLIDINGWINDOW:4:20 MINLEN:70. The version of *A. baumannii* ATCC 17978-mff (GCF_001077675.1 downloaded from the NCBI RefSeq database,17-Mar-2017) was used as the reference genome for variant calling. We added the two additional plasmid sequences present in the *A. baumannii* strain (NC009083, NC_009084) to the chromosome NZ_CP012004 and plasmid NZ_CP012005. Mutations were then manually curated and filtered to remove false positives under the following criteria: mutations were filtered if the gene was found to contain a mutation when the ancestor sequence was compared to the reference genome or if a mutation never reached a cumulative frequency of 10% across all replicate populations.

### Data Availability

R code for filtering and data processing are deposited here: https://github.com/sirmicrobe/U01_allele_freq_code. All sequences were deposited into NCBI under the BioProject number PRJNA485123 and accession numbers can be found in the Supplemental Table S5.

## Supplemental text

### Summary of Travisano et al. experimental design

Following Travisano and coworkers (*1*), consider replicate populations founded by a single clone that are propagated in the same environment for a certain number of generations. We can dissect the roles of each evolutionary force by measuring changes in the mean and variance of an important trait (for example fitness or antibiotic resistance) (Figure 1A). In the first scenario, the mean and variance of the studied trait did not change, so one can conclude that the trait did not evolve (Top left panel, Figure 1A). In the second scenario, while the grand mean of the trait remains the same as the ancestral value, trait variance increases (Top middle panel, Figure 1A). Here, the main evolutionary force is chance, comprised of mutation and genetic drift. In the third scenario, the grand trait mean increases significantly but not the variance (Top right panel, Figure 1A), a change that is best explained by natural selection. Combining these two forces of chance and natural selection, we would expect both trait mean and variance to increase (Bottom left panel, Figure 1A). Note that these four scenarios describe outcomes when starting from a single clone, i.e. with no genetic variation, but this rarely happens in nature. If we conduct the same experiment using three different ancestors that vary in the studied trait, two additional scenarios are possible. In the first, the initial variation among the different ancestors is erased by chance and adaptation (Bottom middle panel, Figure 1A), which cause the trait variance and mean to increase to identical values, regardless of the ancestral value. In the last scenario, the effect of history constrains the evolution of the trait, where the final trait value correlates with the ancestral value (Bottom right panel, Figure 1A) despite contributions of both chance (increased variance) and selection increasing the trait.

**Figure S1.**
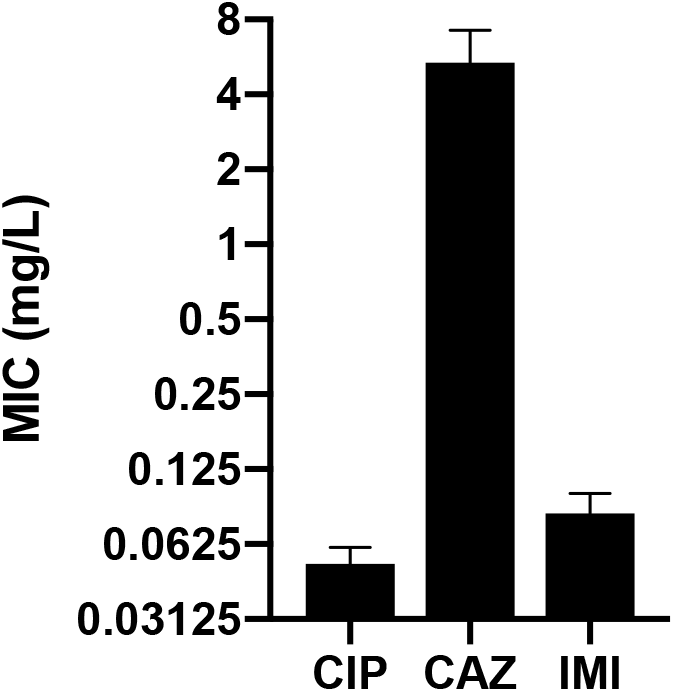
Resistance levels to ciprofloxacin, ceftazidime and imipenem of the ancestral strain prior to being propagated in the historical phase under increasing concentrations of CIP.

**Figure S2.**
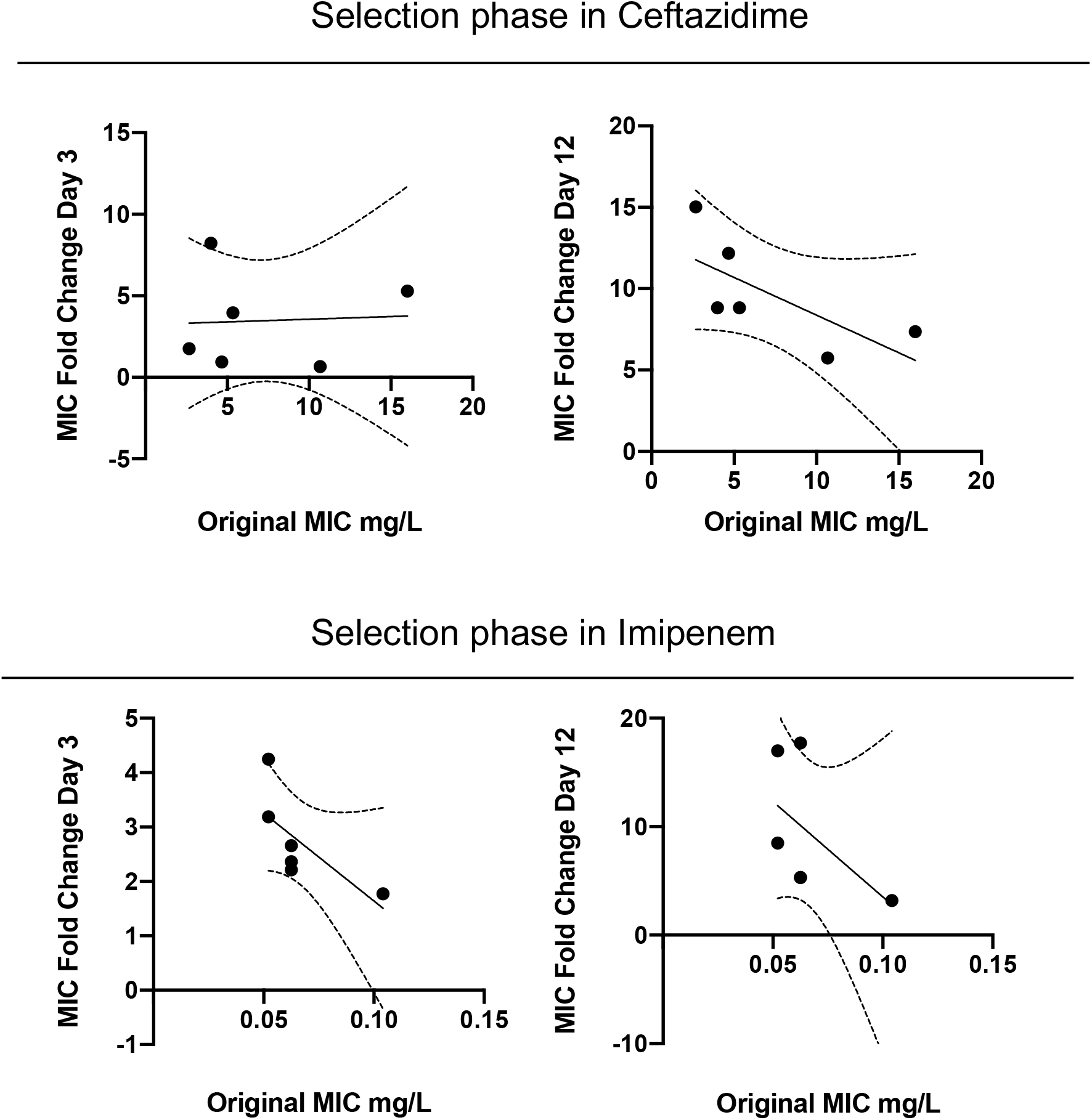
Correlation between ancestral MIC and increase of CAZ (top) and IMI (bottom) resistance after 3 and 12 days evolving in the presence of CAZ (left and right panels respectively). There are three possible outcomes by correlating the original MIC and the fold dilution change: i) a negative correlation, in which the populations with lower initial MICs increased their resistance level more than populations with higher MICs, which implies that the selection erased the previous effects of history; ii) a positive correlation indicates that initial differences in MIC were magnified by selection and iii) a lack of correlation indicates that the effect of history did not change before and after selection. Evolving in presence of CAZ, no correlation was found after three days of antibiotic treatment (Spearman *r* = −0.08, p = 0.919) but the increases in resistance levels and time were negatively correlated with the starting values at day 12 (Spearman *r* = −0.84, p = 0.044) (Figures 2B and S2). Evolving in presence of IMI, increases in resistance levels and time were negatively correlated at day 3 (Spearman *r* = −0.9258, p = 0.033) but not at day 12 (Spearman *r* = −0.6262, p = 0.1833)

**Table S1.**
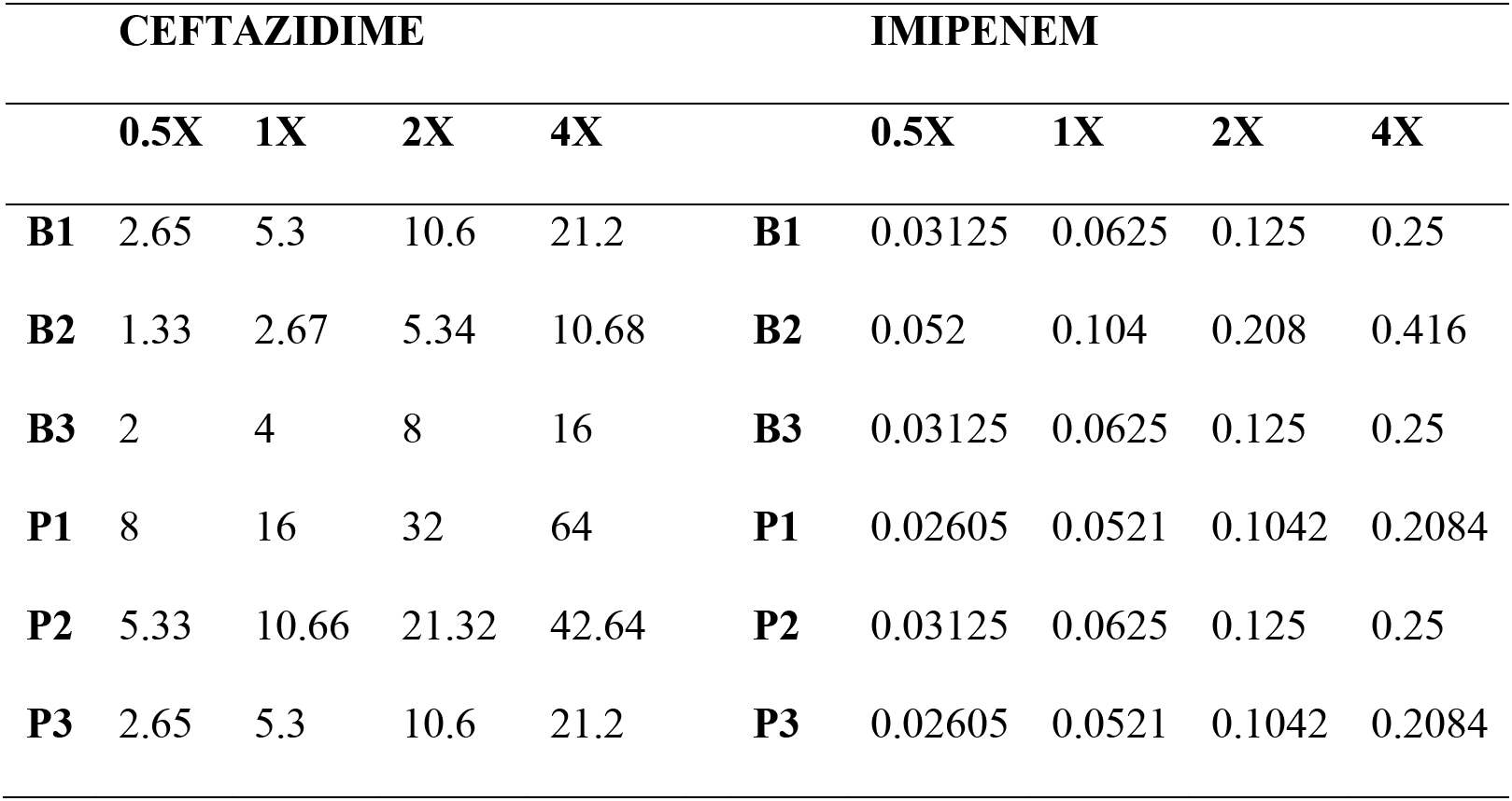
Concentrations of CAZ and IMI (mg/L) added to the broth at different times of the evolution experiments

**Table S2.** Putative driver mutations and resistance levels of the replicate populations after 12 days evolving in presence of CAZ or IMI. The average resistance levels (mg/L) and SEM are shown in the table. Replicates highlighted acquired the same mutation.

| Selection phase in CAZ |  |  |  |  |  |  |  |  |  |
| --- | --- | --- | --- | --- | --- | --- | --- | --- | --- |
|  | Driver mutation in *adeJ* | CAZ (d3) |  | CAZ (d12) |  | IMI (d12) |  | CIP (d12) |  |
| B1_1 | I383F | 21.14 ± | 0.00 | 42.40 ± | 0.00 | 0.33 ± | 0.00 | 0.50 ± | 0.00 |
| B1_2 | *adeN* Δ3bp (541-543) | 21.14 ± | 0.00 | 42.40 ± | 0.00 | 0.12 ± | 0.02 | 0.13 ± | 0.00 |
| B1_3 | Q167R | 21.14 ± | 0.00 | 42.40 ± | 0.00 | 0.17 ± | 0.00 | 1.00 ± | 0.00 |
| B2_1 | Q176K | 2.64 ± | 0.00 | 42.40 ± | 0.00 | 0.17 ± | 0.00 | 0.06 ± | 0.00 |
| B2_2 | K15* | 5.29 ± | 0.00 | 31.80 ± | 6.12 | 0.17 ± | 0.00 | 0.13 ± | 0.00 |
| B2_3 | A290T | 7.93 ± | 1.53 | 42.40 ± | 0.00 | 0.25 ± | 0.05 | 0.13 ± | 0.00 |
| B3_1 | G288S | 21.14 ± | 0.00 | 42.40 ± | 0.00 | 0.17 ± | 0.00 | 0.13 ± | 0.00 |
| B3_2 | F136L | 31.72 ± | 6.10 | 21.20 ± | 0.00 | 0.17 ± | 0.00 | 0.50 ± | 0.00 |
| B3_3 | G288S | 42.29 ± | 0.00 | 42.40 ± | 0.00 | 0.33 ± | 0.00 | 0.13 ± | 0.00 |
| P1_1 | A1007S | 84.57 ± | 0.00 | 84.80 ± | 0.00 | 0.17 ± | 0.00 | 16.03 ± | 0.00 |
| P1_2 | T743P | 84.57 ± | 0.00 | 169.60 ± | 0.00 | 0.33 ± | 0.00 | 16.03 ± | 0.00 |
| P1_3 | Q176K | 84.57 ± | 0.00 | 127.20 ± | 24.48 | 0.17 ± | 0.00 | 8.02 ± | 0.00 |
| P2_1 | Q176R | 10.57 ± | 0.00 | 84.80 ± | 0.00 | 0.08 ± | 0.00 | 8.02 ± | 0.00 |
| P2_2 | *ftsI* Δ3bp (1072-1074) | 5.29 ± | 0.00 | 42.40 ± | 0.00 | 0.08 ± | 0.00 | 0.50 ± | 0.00 |
| P2_3 | Q176R | 7.93 ± | 1.53 | 42.40 ± | 0.00 | 0.17 ± | 0.00 | 8.02 ± | 0.00 |
| P3_1 | V158L (77%), F94L (28%) | 3.96 ± | 0.76 | 63.60 ± | 12.24 | 0.08 ± | 0.00 | 8.02 ± | 0.00 |
| P3_2 | V158L | 2.64 ± | 0.00 | 84.80 ± | 0.00 | 0.17 ± | 0.00 | 8.02 ± | 0.00 |
| P3_3 | F136S | 3.96 ± | 0.76 | 63.60 ± | 12.24 | 0.08 ± | 0.00 | 8.02 ± | 0.00 |
| Selection phase in IMI |  |  |  |  |  |  |  |  |  |
|  | Driver mutation in *ftsI* | IMI (d3) |  | IMI (d12) |  | CAZ (d12) |  | CIP (d12) |  |
| B1_1 | A579V | 0.17 ± | 0.00 | 0.33 ± | 0.00 | 4 ± | 0 | 0.42 ± | 0.07 |
| B1_2 | T506I | 0.14 ± | 0.02 | 0.33 ± | 0.00 | 5.33 ± | 1.09 | 0.50 ± | 0.00 |
| B1_3 | T506I | 0.14 ± | 0.02 | 0.33 ± | 0.00 | 8.00 ± | 0.00 | 0.42 ± | 0.07 |
| B2_1 | A579V | 0.17 ± | 0.00 | 0.33 ± | 0.00 | 4.00 ± | 0.00 | 0.50 ± | 0.00 |
| B2_2 | G524C | 0.22 ± | 0.05 | 0.33 ± | 0.00 | 2.00 ± | 0.00 | 0.42 ± | 0.07 |
| B2_3 | A579T | 0.17 ± | 0.00 | 0.33 ± | 0.00 | 4.00 ± | 0.00 | 0.10 ± | 0.02 |
| B3_1 | A583V | 0.17 ± | 0.00 | 0.17 ± | 0.00 | 4.00 ± | 0.00 | 0.13 ± | 0.00 |
| B3_2 | G574S | 0.08 ± | 0.00 | 0.17 ± | 0.00 | 2.00 ± | 0.00 | 0.08 ± | 0.02 |
| B3_3 | G524S | 0.17 ± | 0.00 | 0.66 ± | 0.00 | 2.00 ± | 0.00 | 0.06 ± | 0.00 |
| P1_1 | H530Y | 0.33 ± | 0.00 | 0.33 ± | 0.00 | 16.00 ± | 0.00 | 16.03 ± | 0.00 |
| P1_2 | A583D | 0.17 ± | 0.00 | 0.33 ± | 0.00 | 10.67 ± | 2.18 | 10.69 ± | 2.18 |
| P1_3 | A579V | 0.17 ± | 0.00 | 0.66 ± | 0.00 | 5.33 ± | 1.09 | 16.03 ± | 0.00 |
| P2_1 | S395V | 0.17 ± | 0.00 | 1.33 ± | 0.00 | 16.00 ± | 0.00 | 2.67 ± | 1.09 |
| P2_2 | A579V | 0.17 ± | 0.00 | 1.33 ± | 0.00 | 53.33 ± | 8.71 | 4.01 ± | 0.00 |
| P2_3 | P580A | 0.17 ± | 0.00 | 0.66 ± | 0.00 | 10.67 ± | 2.18 | 5.34 ± | 1.09 |
| P3_1 | S539P | 0.17 ± | 0.00 | 1.33 ± | 0.00 | 8.00 ± | 0.00 | 4.01 ± | 0.00 |
| P3_2 | A578T | 0.17 ± | 0.00 | 0.66 ± | 0.00 | 4.00 ± | 0.00 | 8.02 ± | 0.00 |
| P3_3 | A579V | 0.17 ± | 0.00 | 0.66 ± | 0.00 | 2.00 ± | 0.00 | 8.02 ± | 0.00 |

**Table S3. Complete list of mutated genes from the sequenced populations and clones.** Complete list of mutated genes obtained and analyzed obtained as explained in methods.

**Table S4. Estimated statistics for history, chance and selection forces.** By means of a nested linear mixed model, the estimated coefficients representing the forces are shown, in addition to the confidence intervals at a α=0.05 significance level generated by bootstrapping and the Bayes Factors computed by a Bayesian analysis. BF_10_ is the ratio between the probabilities of the alternative and null model and therefore, it measures the degree of evidence of including the force. BF_10_ < 1 null evidence, 3 > BF_10_ > 1 weak evidence, 20 > BF_10_ > 3 positive evidence, 150 > BF_10_ > 20 strong evidence, BF_10_ > 150 very strong evidence.

**Table S5. List of deposited sequences from clones and populations and the corresponding accession numbers.**

## References

1. M. Travisano, J. A. Mongold, A. F. Bennett, R. E. Lenski, Experimental tests of the roles of adaptation, chance, and history in evolution. Science. 267, 87–90 (1995).

2. M. Rebolleda-Gomez, M. Travisano, Adaptation, chance, and history in experimental evolution reversals to unicellularity. Evolution. 73, 73–83 (2019).

3. J. R. Meyer, D. T. Dobias, J. S. Weitz, J. E. Barrick, R. T. Quick, R. E. Lenski, Repeatability and contingency in the evolution of a key innovation in phage lambda. Science. 335, 428–32 (2012).

4. S. Kryazhimskiy, D. P. Rice, E. R. Jerison, M. M. Desai, Global epistasis makes adaptation predictable despite sequence-level stochasticity. Science. 344, 1519–1522 (2014).

5. S. R. Keller, D. R. Taylor, History, chance and adaptation during biological invasion: separating stochastic phenotypic evolution from response to selection. Ecol Lett. 11, 852–66 (2008).

6. Z. D. Blount, R. E. Lenski, J. B. Losos, Contingency and determinism in evolution: Replaying life’s tape. Science. 362 (2018), doi:10.1126/science.aam5979.

7. M. Lassig, V. Mustonen, A. M. Walczak, Predicting evolution. Nature Ecology & Evolution. 1, 77 (2017).

8. C. B. Turner, C. W. Marshall, V. S. Cooper, Parallel genetic adaptation across environments differing in mode of growth or resource availability. Evolution Letters. 2, 355–367 (2018).

9. S. F. Bailey, N. Rodrigue, R. Kassen, The effect of selection environment on the probability of parallel evolution. Molecular Biology and Evolution. 32, 1436–48 (2015).

10. M. Galardini, B. P. Busby, C. Vieitez, A. S. Dunham, A. Typas, P. Beltrao, The impact of the genetic background on gene deletion phenotypes in Saccharomyces cerevisiae. Molecular Systems Biology. 15, e8831 (2019).

11. Z. D. Blount, C. Z. Borland, R. E. Lenski, Historical contingency and the evolution of a key innovation in an experimental population of <em>Escherichia coli</em>. Proceedings of the National Academy of Sciences. 105, 7899–7906 (2008).

12. M. L. Salverda, E. Dellus, F. A. Gorter, A. J. Debets, J. van der Oost, R. F. Hoekstra, D. S. Tawfik, J. A. de Visser, Initial mutations direct alternative pathways of protein evolution. PLoS Genetics. 7, e1001321 (2011).

13. A. I. Khan, D. M. Dinh, D. Schneider, R. E. Lenski, T. F. Cooper, Negative epistasis between beneficial mutations in an evolving bacterial population. Science. 332, 1193–6 (2011).

14. T. Vogwill, M. Kojadinovic, V. Furio, R. C. MacLean, Testing the role of genetic background in parallel evolution using the comparative experimental evolution of antibiotic resistance. Molecular Biology and Evolution. 31, 3314–23 (2014).

15. M. R. Scribner, A. Santos-Lopez, C. W. Marshall, C. Deitrick, V. S. Cooper, Parallel Evolution of Tobramycin Resistance across Species and Environments. mBio. 11 (2020), doi:10.1128/mBio.00932-20.

16. E. Wistrand-Yuen, M. Knopp, K. Hjort, S. Koskiniemi, O. G. Berg, D. I. Andersson, Evolution of high-level resistance during low-level antibiotic exposure. Nat Commun. 9, 1599 (2018).

17. A. R. Hall, R. C. MacLean, Epistasis buffers the fitness effects of rifampicin- resistance mutations in Pseudomonas aeruginosa. Evolution. 65, 2370–9 (2011).

18. D. R. Gifford, R. Krasovec, E. Aston, R. V. Belavkin, A. Channon, C. G. Knight, Environmental pleiotropy and demographic history direct adaptation under antibiotic selection. Heredity. 121, 438–448 (2018).

19. P. Yen, J. A. Papin, History of antibiotic adaptation influences microbial evolutionary dynamics during subsequent treatment. PLoS Biol. 15, e2001586 (2017).

20. S. Trindade, A. Sousa, K. B. Xavier, F. Dionisio, M. G. Ferreira, I. Gordo, Positive epistasis drives the acquisition of multidrug resistance. PLoS Genetics. 5, e1000578 (2009).

21. A. Santos-Lopez, C. W. Marshall, M. R. Scribner, D. J. Snyder, V. S. Cooper, Evolutionary pathways to antibiotic resistance are dependent upon environmental structure and bacterial lifestyle. eLife. 8, e47612 (2019).

22. V. S. Cooper, Experimental Evolution as a High-Throughput Screen for Genetic Adaptations. mSphere. 3 (2018), doi:10.1128/mSphere.00121-18.

23. B. H. Good, M. J. McDonald, J. E. Barrick, R. E. Lenski, M. M. Desai, The dynamics of molecular evolution over 60,000 generations. Nature. 551, 45–50 (2017).

24. A. N. Nguyen Ba, I. Cvijović, J. I. Rojas Echenique, K. R. Lawrence, A. Rego-Costa, X. Liu, S. F. Levy, M. M. Desai, High-resolution lineage tracking reveals travelling wave of adaptation in laboratory yeast. Nature. 575, 494–499 (2019).

25. CDC, The biggest antibiotic-resistant threats in the U.S. Centers for Disease Control and Prevention (2019), (available at https://www.cdc.gov/drugresistance/biggest-threats.html).

26. R. C. MacLean, A. San Millan, The evolution of antibiotic resistance. Science. 365, 1082–1083 (2019).

27. R. Pokhriyal, R. Hariprasad, L. Kumar, G. Hariprasad, Chemotherapy Resistance in Advanced Ovarian Cancer Patients. Biomarkers in cancer. 11, 1179299x19860815 (2019).

28. D. Hughes, D. I. Andersson, Evolutionary consequences of drug resistance: shared principles across diverse targets and organisms. Nat Rev Genet. 16, 459–71 (2015).

29. B. K. Verlinden, A. Louw, L. M. Birkholtz, Resisting resistance: is there a solution for malaria? Expert Opin Drug Discov. 11, 395–406 (2016).

30. M. Palaci, R. Dietze, D. J. Hadad, F. K. C. Ribeiro, R. L. Peres, S. A. Vinhas, E. L. N. Maciel, V. do Valle Dettoni, L. Horter, W. H. Boom, J. L. Johnson, K. D. Eisenach, Cavitary Disease and Quantitative Sputum Bacillary Load in Cases of Pulmonary Tuberculosis. J Clin Microbiol. 45, 4064–4066 (2007).

31. I. K. Paterson, A. Hoyle, G. Ochoa, C. Baker-Austin, N. G. H. Taylor, Optimising Antibiotic Usage to Treat Bacterial Infections. Scientific Reports. 6, 37853 (2016).

32. R. M. Jones, M. Nicas, A. Hubbard, M. D. Sylvester, A. Reingold, The Infectious Dose of Francisella Tularensis (Tularemia). Appl Biosaf. 10, 227–239 (2005).

33. M. J. Wiser, N. Ribeck, R. E. Lenski, Long-Term Dynamics of Adaptation in Asexual Populations. Science. 342, 1364–1367 (2013).

34. C. Pal, B. Papp, V. Lazar, Collateral sensitivity of antibiotic-resistant microbes. Trends in Microbiology. 23, 401–7 (2015).

35. C.-R. Lee, J. H. Lee, M. Park, K. S. Park, I. K. Bae, Y. B. Kim, C.-J. Cha, B. C. Jeong, S. H. Lee, Biology of Acinetobacter baumannii: Pathogenesis, Antibiotic Resistance Mechanisms, and Prospective Treatment Options. Front Cell Infect Microbiol. 7, 55 (2017).

36. S. Bedhomme, G. Lafforgue, S. F. Elena, Genotypic but not phenotypic historical contingency revealed by viral experimental evolution. BMC Evolutionary Biology. 13, 46 (2013).

37. A. Dunai, R. Spohn, Z. Farkas, V. Lázár, Á. Györkei, G. Apjok, G. Boross, B. Szappanos, G. Grézal, A. Faragó, L. Bodai, B. Papp, C. Pál, Rapid decline of bacterial drug-resistance in an antibiotic-free environment through phenotypic reversion. eLife. 8, e47088 (2019).

38. P. Durão, R. Balbontín, I. Gordo, Evolutionary Mechanisms Shaping the Maintenance of Antibiotic Resistance. Trends in Microbiology. 26, 677–691 (2018).

39. H. Teotonio, M. R. Rose, Variation in the reversibility of evolution. Nature. 408, 463–6 (2000).

40. E. Kugelberg, S. Lofmark, B. Wretlind, D. I. Andersson, Reduction of the fitness burden of quinolone resistance in Pseudomonas aeruginosa. J Antimicrob Chemother. 55, 22–30 (2005).

41. S. J. Gould, Wonderful Life: The Burgess Shale and the Nature of History (W. W. Norton & Company, 1990).

42. W. Loftie-Eaton, K. Bashford, H. Quinn, K. Dong, J. Millstein, S. Hunter, M. K. Thomason, H. Merrikh, J. M. Ponciano, E. M. Top, Compensatory mutations improve general permissiveness to antibiotic resistance plasmids. Nature Ecology & Evolution (2017), doi:10.1038/s41559-017-0243-2.

43. S. J. Dunn, C. Connor, A. McNally, The evolution and transmission of multi-drug resistant Escherichia coli and Klebsiella pneumoniae: the complexity of clones and plasmids. Current Opinion in Microbiology. 51, 51–56 (2019).

44. M. Baym, S. Kryazhimskiy, T. D. Lieberman, H. Chung, M. M. Desai, R. Kishony, Inexpensive multiplexed library preparation for megabase-sized genomes. PLoS One. 10, e0128036 (2015).

45. E.-J. Wagenmakers, A practical solution to the pervasive problems ofp values. Psychonomic Bulletin & Review. 14, 779–804 (2007).

46. J. E. Barrick, G. Colburn, D. E. Deatherage, C. C. Traverse, M. D. Strand, J. J. Borges, D. B. Knoester, A. Reba, A. G. Meyer, Identifying structural variation in haploid microbial genomes from short-read resequencing data using breseq. BMC Genomics. 15, 1039 (2014).

47. A. M. Bolger, M. Lohse, B. Usadel, Trimmomatic: a flexible trimmer for Illumina sequence data. Bioinformatics. 30, 2114–2120 (2014).

